# Information gathering explains decision dynamics during human and monkey reward foraging

**DOI:** 10.1101/2023.10.14.562362

**Authors:** David L Barack, Felipe Parodi, Vera Ludwig, Michael L Platt

## Abstract

Foraging in humans and other animals requires a delicate balance between exploitation of current resources and exploration for new ones. The tendency to overharvest—lingering too long in depleting patches—is a routine behavioral deviation from predictions of optimal foraging theories. To characterize the computational mechanisms driving these deviations, we modeled foraging behavior using a virtual patch-leaving task with human participants and validated our findings in an analogous foraging task in two monkeys. Both humans and monkeys overharvested and stayed longer in patches with longer travel times compared to shorter ones. Critically, patch residence times in both species declined over the course of sessions, enhancing reward rates in humans. These decisions were best explained by a logistic transformation that integrated both current rewards and information about declining rewards. This parsimonious model demystifies both the occurrence and dynamics of overharvesting, highlighting the role of information gathering in foraging. Our findings provide insight into computational mechanisms shaped by ubiquitous foraging dilemmas, underscoring how behavioral modeling can reveal underlying motivations of seemingly irrational decisions.

## 1. Introduction

Foraging for resources is a fundamental behavior in species across the phylogenetic spectrum. Foraging involves iterated accept-or-reject decisions about persistent resources rather than one-shot decisions (Barack and Platt 2017; Hayden 2018). Such foraging choice contexts are ubiquitous, in both more traditional contexts, such as Amazonian hunting (Hill et al. 1987), Nahua mushroom gathering (Pacheco-Cobos et al. 2019), or Inuit ungulate hunting (Hall Jr 1971; Condon et al. 1995), and more urban ones like shopping (Rajala and Hantula 2000; Foxall and James 2003; Hantula et al. 2008). Foraging is increasingly utilized to characterize the search for other resources beyond food, such as through the space of concepts (Hills et al. 2010), memories (Hills et al. 2012), or strategies (Metcalfe and Kornell 2005; Metcalfe and Jacobs 2010). Further, signatures of foraging have been identified across diverse tasks, ranging from surfing on the internet (Fu and Pirolli 2007) to searching for shapes hidden on a grid (Barack et al. 2023).

Given the breadth and evolutionary significance of foraging, a general characterization of how humans and other animals make foraging decisions may provide insights into aspects of human decision making that are difficult to explain from the perspective of neo-classical economics (Gigerenzer and Gaissmaier 2011; Barack and Platt 2017; Hayden 2018). Foraging can be dissected into decisions about when to leave a depleting resource and decisions about where to search next (Hills et al. 2010). Here, we characterize decisions to leave resources by studying both human and monkey participants engaged in virtual foraging tasks in order to model the basic computations and representations supporting these decisions. In our task, participants made repeated decisions either to exploit and continue harvesting depleting resources or to explore by leaving the depleting resource for a new undepleted one. This choice context mimics both the depletion and renewal of patchy concentrations of resources across spatial and temporal domains, ubiquitous in natural environments (Levin 1994), as well as decisions to explore or exploit depleting resources that often go awry in psychiatric disorders (Addicott et al. 2017). By modeling the computations underlying humans’ and monkeys’ performance, we can gain insight into the mechanisms governing this evolutionarily significant class of behaviors.

Optimal foraging theory provides a body of formal models of exploratory decisions in well-circumscribed cases (Charnov 1976; Stephens and Krebs 1986). Under certain assumptions, the Marginal Value Theorem states that when the instantaneous intake rate drops below the average for the environment, foragers should depart the current patch (Charnov 1976). In stochastic environments, the Potential Value Theorem states that foragers should stay when the potential value of a patch is positive and leave when it drops to zero (McNamara 1982). More recently, Bayesian foraging theory has expanded the scope of foraging theory to explain purported suboptimalities in patch-leaving behavior including overharvesting (Houston and McNamara 1982; Olsson and Holmgren 1998; Davidson and El Hady 2019; Kilpatrick et al. 2021).

While foraging animals tend to qualitatively obey the predictions of optimal foraging theory by balancing current offers against long-term expectations, they sometimes deviate from the quantitative predictions of these models. Foragers often overharvest, persisting in patches beyond the optimal leave time (Nonacs 2001; Hayden et al. 2011; Kolling et al. 2012; Wikenheiser et al. 2013; Shenhav et al. 2014; Constantino and Daw 2015; Carter and Redish 2016; Kane et al. 2019; Kendall and Wikenheiser 2022). Various explanations for this deviation have been proposed, including aversions to rejecting immediately available rewards (Wikenheiser et al. 2013; Carter and Redish 2016), nonlinear utility functions (Constantino and Daw 2015), intertemporal choice preferences (such as short-term rate maximization (Stephens 2002; Stephens et al. 2004; Namboodiri et al. 2014) or intertemporal discounting (Ainslie and Haslam 1992; Kirby 1997; Kane et al. 2019)), and gathering evidence about the environment (Davidson and El Hady 2019; Harhen and Bornstein 2023). Notably, foragers not only tend to overstay in patches but this overharvesting is dynamic, changing as animals have more experience with the environment (Barack et al. 2022; Harhen and Bornstein 2023). These dynamics may provide novel insights into the computational processes supporting foraging decisions.

To address the dynamics of these deviations from predicted optimal behavior, we studied human participants making repeated explore-exploit foraging decisions and fit numerous models to their behavior. We extended our findings by fitting the same set of models to foraging decisions made by rhesus macaques. Consistent with prior findings, both human and monkey participants were sensitive to the costs of exploration but also stayed longer in depleting patches than predicted by optimal foraging models. Total time spent foraging in depleting patches declined over the course of sessions, a potential clue to the computations used to make patch leaving decisions. For both humans and monkeys, the best fit model used a mixture of currencies, including reward and information, and computed decisions to leave using a simple logistic transformation. This model can explain both overharvesting and its dynamics, and predicts observed decreases in patch residence times as foragers accumulated experience within the environment. These findings indicate that humans and monkeys gather both rewards and information while foraging and combine them to determine when to leave a depleting patch and search for a new one.

## 2. Results

### Online Patch Leaving Task and Behavior

Adults (n = 422; see methods) performed an online foraging task (Fig 1; data used for analysis and modeling were first reported in (Barack et al. 2022)). Participants were recruited via Prolific (54.59% male, average age = 44.66 ± 15.98 y) and were instructed to collect as many berries as possible in 8 min (4 blocks, 2 min each). On each trial, participants chose between staying at a given patch, in which payoffs declined with each successive choice, and traveling to a new patch, which took time. Each participant was tested in two conditions that differed only in the timeout to reset bushes to the full value (either 1 sec or 5 sec), indicated by the visual distance between bushes. The order of conditions (either 1 sec - 5 sec - 1 sec - 5 sec or 5 - 1 - 5 - 1) was counterbalanced across participants.

**Figure 1.**
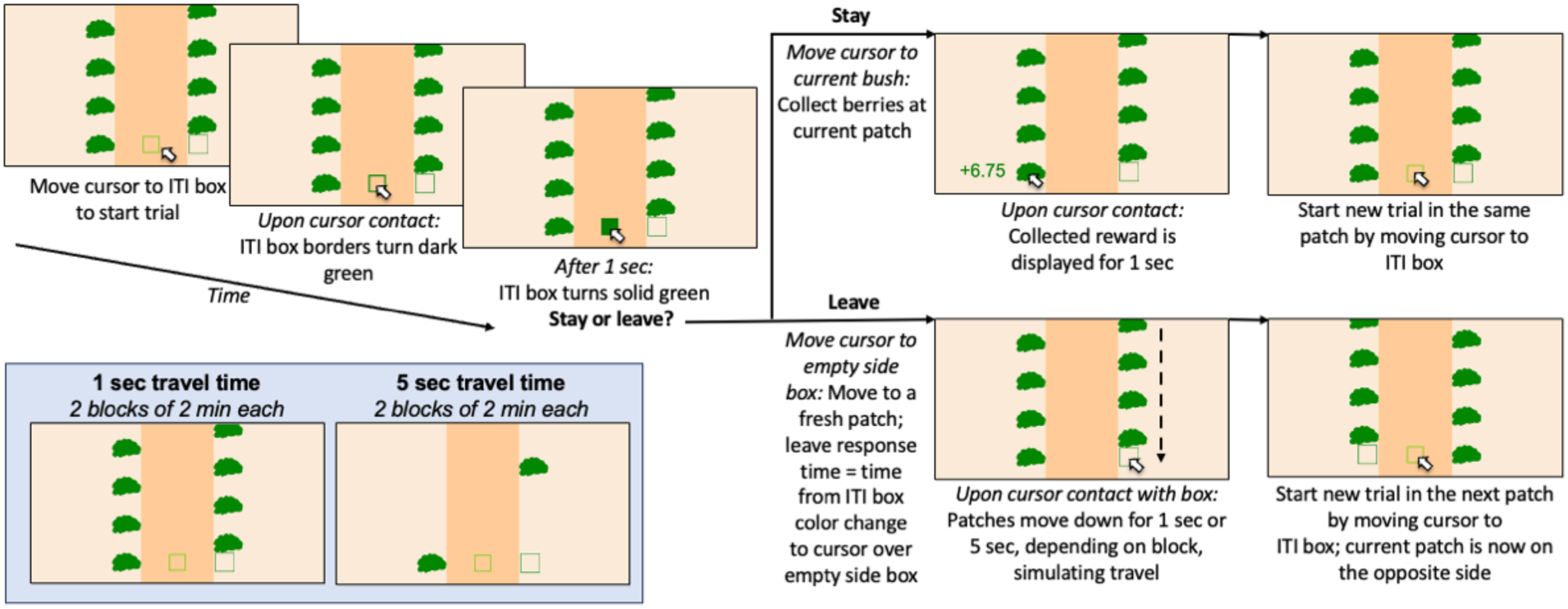
Virtual Foraging Task. On each trial, participants began by moving a cursor to an intertrial interval (ITI) box. After 1 sec, the ITI box turned solid green. Participants then chose between staying at a given patch to gather points (exploit decision), the expected payoffs of which declined over time, or replenishing the patch in exchange for a timeout delay (explore decision). Timeouts were either 1 sec or 5 sec, in blocks. Inset: screenshots of the two foraging environments, 1 sec timeouts on the left and 5 sec on the right.

Participants left patches in short timeout (1 sec) environments earlier than in long timeout (5 sec) ones, as predicted by foraging theory (paired t-test on median patch residence time (time in patch *TiP*) by participant, t(df = 421) = -12.13, p < 1×10^-28^; 1 sec timeout *TiP*_short_ = 16.32 ± 0.47 sec; 5 sec timeout *TiP*_long_ = 20.99 ± 0.51 sec). Participants earned significantly more points on average in long timeout environments than short ones (paired t-test on median reward by patch and timeout, t(df = 421) = -14.98, p < 1×10^- 40^; *R*_short_ = 24.76 ± 0.60 pts; *R*_long_ = 31.94 ± 0.61 pts), but did not harvest rewards at significantly different rates (paired t-test on median reward rate by patch and timeout, p > 0.25, t(df = 421) = -1.11; reward rate *RR*_short_ = 1.54 ± 0.030 pts/sec; *RR*_long_ = 1.56 ± 0.025 pts/sec). Average reward rates were significantly less than the optimal reward rates in both short timeout environments (*RR*_short,optimal_ = 2.61 points/sec; one-sample t-test, t(df = 421) = -36.07, p < 1×10^-130^) and long timeout environments (*RR*_long,optimal_ = 2.23, p < 1×10^-75^, t(df = 421) = -23.02). This suboptimal behavior resulted from participants staying too long in each patch (Figure 2; one-sample t-tests; short patches: *TiP*_short,optimal_ = 6.06 sec, t(df = 421) = 21.66, p < 1×10^-69^; long patches: *TiP*_long,optimal_ = 12.90 sec, p < 1×10^-44^, t(df = 421) = 15.89), as observed in prior studies in other animals including monkeys (Pyke et al. 1977; Nonacs 2001; Hayden et al. 2011; Kendall and Wikenheiser 2022).

**Figure 2.**
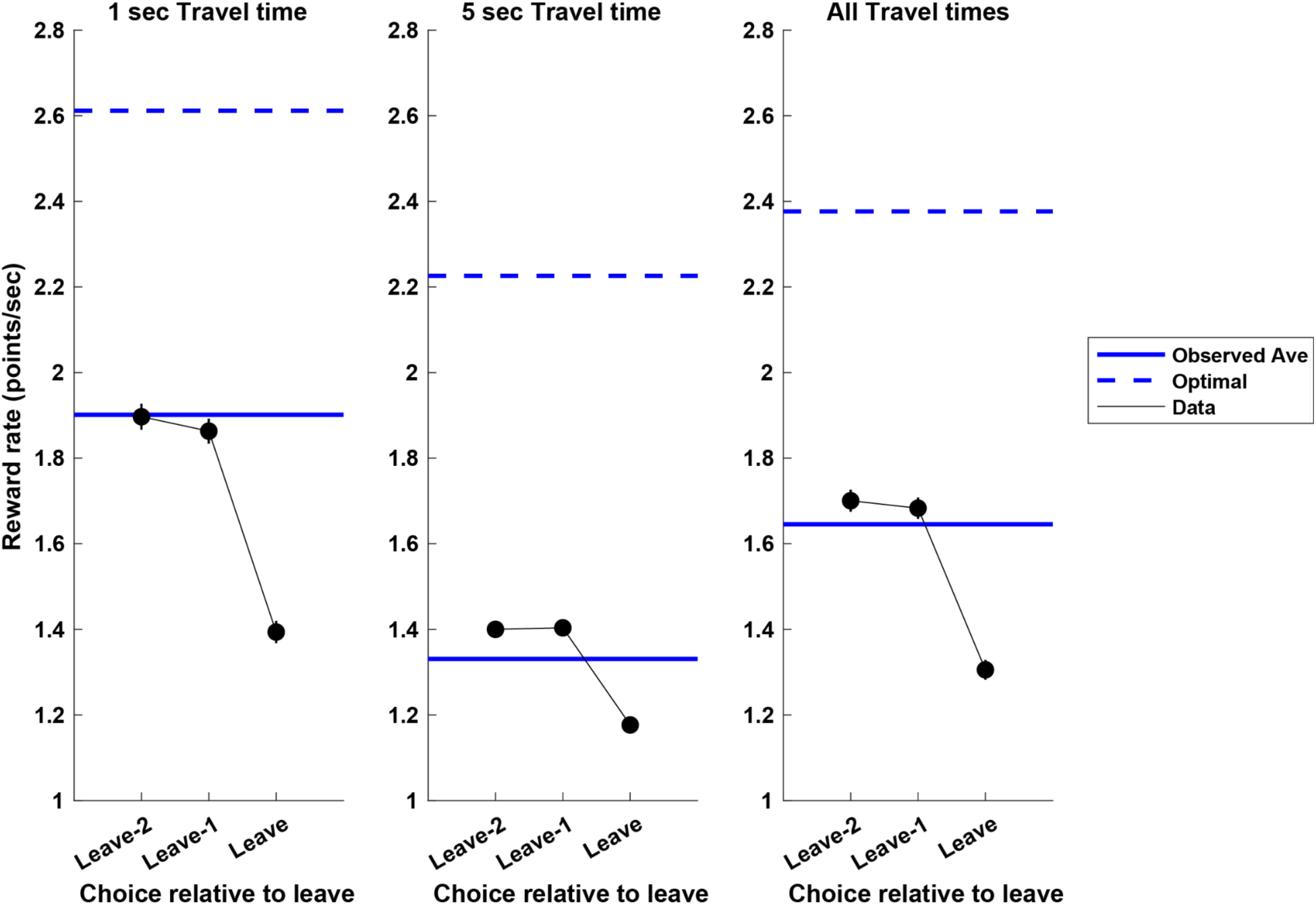
Humans overharvest during patch foraging. Human participants (n = 425) overharvested patches relative to numerically computed optimal reward rates. X-axis: choice relative to a decision to leave; y-axis: reward rate; black points: mean ± s.e.m.; solid blue line: observed average reward rate across all choices in patch; dashed blue line: reward rates at which leaving a patch is optimal, i.e., maximizes long-term reward intake (see methods). Error bars are sometimes hidden by data points.

Patch harvesting varied both across participants (Fig. 3A) and blocks (Fig. 3B) during the session, consistent with recent reports (Barack et al. 2022; Harhen and Bornstein 2023). Participants performed up to four blocks of patches, alternating between 1 sec and 5 sec (or 5 sec and 1 sec) timeout blocks (see methods). The mean of participants’ median patch residence times were significantly higher for the first compared to the second block for the longer 5 sec delay condition (paired t-test, t(df=415) = 3.45, p < 0.001; first block mean median *TiP*_long,1_ = 24.46 ± 0.81 sec; second block *TiP*_long,2_ = 21.78 ± 0.55 sec; Fig. 3B right), though not for the shorter 1 sec delay (paired t- test, t(df=416) = 0.55, p > 0.55; *TiP*_short,1_ = 17.91 ± 0.60 sec; *TiP*_short,2_ = 17.51 ± 0.58 sec; Fig. 3B left). After concatenating together the two blocks for each timeout, some participants showed an effect of patch number in session on patch residence time, typically decreasing residence times across patches (1 sec patches: OLS, 111 / 425 participants significant (p < 0.05) slope, 74 negative; 5 sec patches: OLS, 85 / 425 participants significant slope (p < 0.05), 53 negative; 166 / 422 unique participants showed some effect). This effect was significant for all patches across the population for both short (Student’s t-test, p < 0.005, t(421) = -2.97, mean β_slope_ = -0.42 ± 0.14) and long timeout blocks (Student’s t-test, p < 0.001, t(421) = -3.35, mean β_slope_ = -0.79 ± 0.24). Data from two example participants are shown in Figure 4. Participant 5 (Figure 4A) did not show an effect of patch number in block (1 sec patches: OLS, β_slope_ = -0.10 ± 0.21 sec/patch_number, p > 0.64; 5 sec patches: OLS, β_slope_ = 0.38 ± 0.44 sec/patch_number, p > 0.42), whereas participant 251 (Figure 4B) did (1 sec patches: OLS, β_slope_ = -0.86 ± 0.15 sec/patch_number, p < 0.001; 5 sec patches: OLS, β_slope_ = -1.58 ± 0.27 sec/patch_number, p < 0.001).

**Figure 3.**
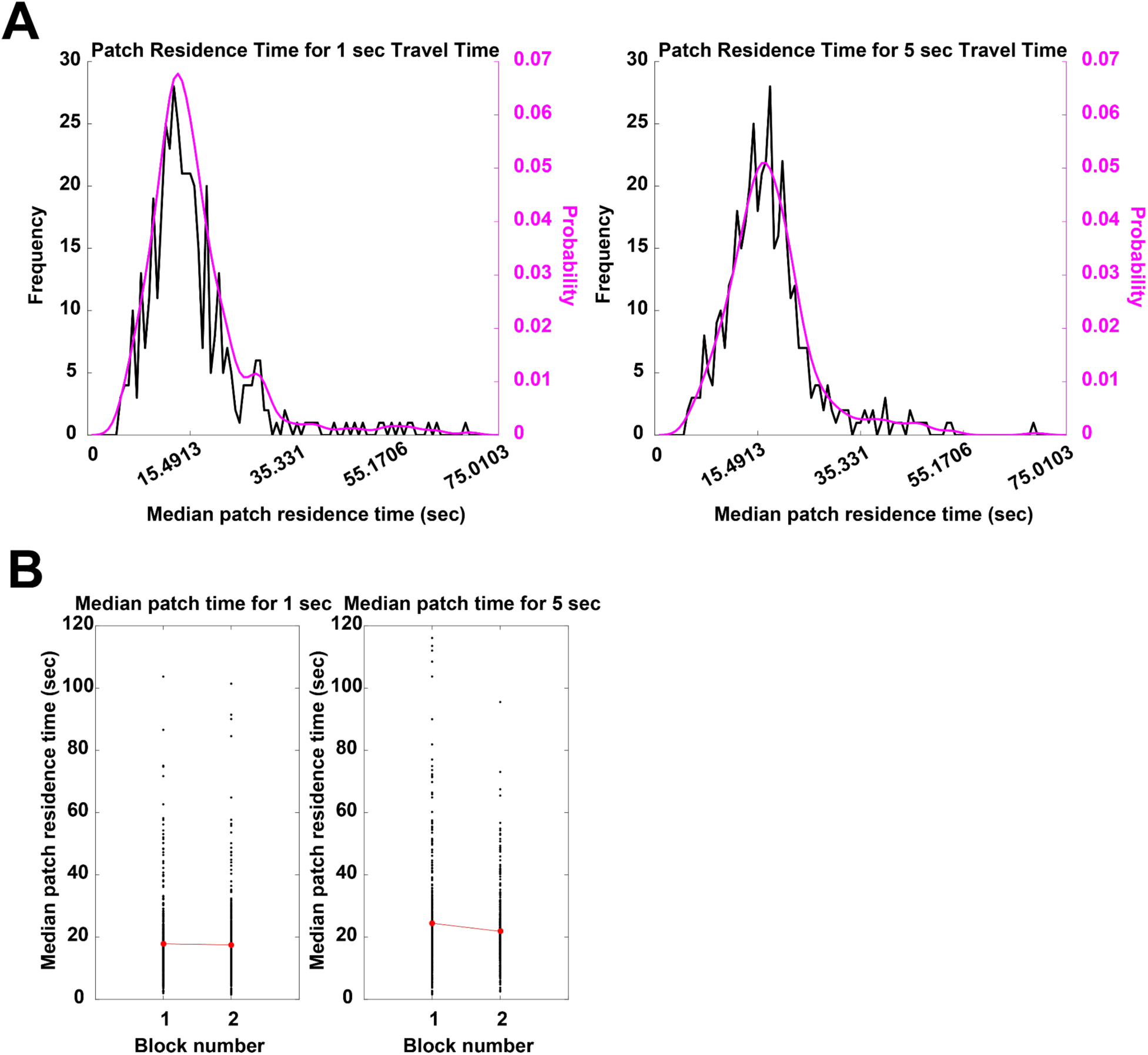
Patch residence times changed over the duration of the experiment. **A.** Human participants (n = 422) showed a distribution of patch harvest times for both 1 sec (left) and 5 sec (right) delays to reset patches to their full value. Black: binned median patch residence time; magenta: kernel density estimates of patch residence time distributions. **B.** Mean median patch residence times for the second block of 5 sec delay patches were significantly shorter than for the first block (right), though no difference was observed across blocks for the 1 sec delay. Black points: individual median patch residence times by delay and block number; red points: mean median patch residence times. Error bars hidden by data points.

**Figure 4.**
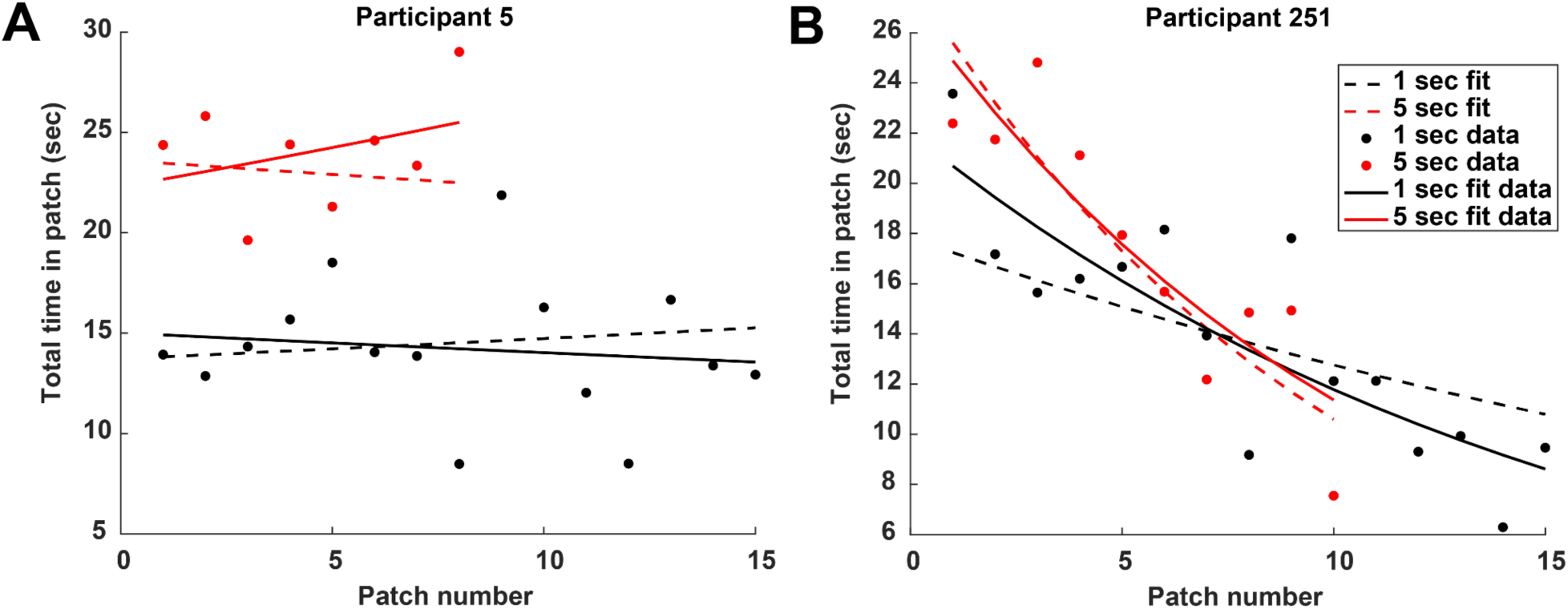
Human participants show either a shallow or a steep decline in patch residence time as a function of experience. Figure depicts the observed change in patch residence time in both 1 sec timeout and 5 sec timeout patches as a function of patch number in concatenated blocks (solid lines) compared to simulations of the best-fit mixture-of- currencies model (dashed lines). On the y-axis is the total time spent gathering points in patch, and on the x-axis is the patch number in session. Points are observed patch residence times. Solid curves are exponential fits to the data; dashed curves are exponential fits to simulated residence times from the best-fit model (1000 iterations; individual simulation data points not shown). A. Participant no. 5, who did not show a patch number in concatenated block effect. B. Participant no. 251, who showed the effect.

Some participants also showed an effect of patch number in concatenated blocks on reward rate, typically increasing reward rates across patches in concatenated blocks (1 sec patches: OLS, 132 / 422 participants significant (p < 0.05) slope, 120 positive; 5 sec patches: OLS, 95 / 425 participants significant slope, 77 positive; 182 / 422 unique participants showed some effect). This effect was significant across the population for both concatenated short timeout blocks (Student’s t-test, p < 1×10^-17^, t(421) = 9.04, mean β_slope_ = 0.038 ± 0.0042) and concatenated long timeout blocks (Student’s t-test, p < 1×10^-14^, t(421) = 8.21, mean β_slope_ = 0.043 ± 0.0052). The same two participants who showed an effect of patch number in concatenated block on patch residence time also illustrate these effects on reward rate, with participant 5 showing no effect of patch number in block on reward rate (1 sec patches: OLS, β_slope_ = 0.0031 ± 0.014 (berries/sec)/patch_number, p > 0.83; 5 sec patches: OLS, β_slope_ = 0.012 ± 0.028 (berries/sec)/patch_number, p > 0.68), unlike participant 251 who did (1 sec patches: OLS, β_slope_ = 0.055 ± 0.011 (berries/sec)/patch_number, p < 0.001; 5 sec patches: OLS, β_slope_ = 0.13 ± 0.017 (berries/sec)/patch_number, p < 0.001). Many participants showed increasing reward rates over patches within concatenated blocks, suggesting learning as a function of experience. As patch residence times declined and reward rates increased across the session, participants enhanced their foraging performance. Hence, participants’ decisions to leave patches sooner and sooner over the duration of the experiment improved performance. These decision dynamics are critical, suggesting participants continuously updated computations governing patch departure as they accumulated experience about the local foraging environment.

### Computations for Patch Leaving Decisions during Foraging

In order to infer the computations that underlie foraging decisions on our task, we fit 9 models to each participant’s behavior. These models can be classified as reinforcement learning, optimal foraging theory, and structural learning models. Reinforcement learning models (Rescorla and Wagner 1972; Sutton and Barto 1998) propose that foragers keep a cached value of the explore and exploit options, learn these cached values over time, and on each trial use that cached value to make a decision to stay or leave a patch. Optimal foraging theory (Stephens and Krebs 1986), a branch of formal ecology, explains foraging behavior in terms of mathematical models, specifically the marginal value theorem (MVT, Charnov 1976), which predicts patch leaving decisions on the basis of a comparison of instantaneous resource intake to average intakes. The MVT instructs foragers to depart depleting patches when the instantaneous intakes fall below that average. Structural learning models (Gershman and Niv 2010; Harhen and Bornstein 2023) explain foraging behavior as the result of both the drive to find reward as well as to learn the structure of the environment. These three model classes were chosen because of their widespread use, their computational interpretability, or their theoretical utility given the nature of our task. Finally, we added a mixture-of-currencies logistic model, which included an information currency in addition to the foraging currencies we consider (discussed below). We reasoned that participants not only gather rewards but also information throughout the tasks, and that this information may impact decisions to leave patches. In order to simplify the interpretation of any role for information in these decisions, we elected to use a simple logistic transformation to predict patch leaving decisions using the integrated value of reward and information. (See methods for longer descriptions of each model.)

We fit each model to behavior using three different currencies (reward, time, and reward rate), yielding a total of 27 models. The first currency was reward, the recently received number of points for staying in a patch. The second currency was time, the cumulative time spent gathering points in a patch. The third currency was reward rate; for some models, reward rate was the number of points gathered in a patch divided by the time in patch, whereas for others, the *instantaneous* reward rate was the relevant currency, set to the expected number of points divided by the sum of the handling time and task-imposed delays (see methods). These currencies were selected due to their relevance in a range of theoretical and empirical studies of foraging (Charnov 1976; McNamara and Houston 1986; Stephens and Krebs 1986; Bateson and Kacelnik 1996; Mcnamara and Houston 1997; Davidson and El Hady 2019; Kane et al. 2019).

In addition to these three currencies, in our mixture-of-currencies model we used a fourth currency, the information gained by observing the decrease in reward from deciding to stay in patch. The reward gathered from staying in patch dropped by an amount drawn from a distribution with mean = 0.5 and standard deviation = 0.25. Participants were not informed of the size and variance in this drop; rather, they had to learn about this distribution by repeatedly experiencing the decrease in reward from staying in a patch. The reward received after each decision to stay in a patch provided evidence about this distribution. We formalized this information as the change in entropy of a normal distribution of experienced drops in reward before and after a new decision to stay in a patch (see methods). This information was then included in our mixture-of- currencies logistic model.

To determine the best-fit model, we selected the model with the best AIC score and computed the protected exceedance probabilities (Stephan et al. 2009; Rigoux et al. 2014). Findings were validated using BIC (see methods). The best fit model was the mixture-of-currencies logistic model (see Table 1). This model contained three value variables for 1 sec and 5 sec timeouts separately: reward rate, information, and their interaction. Hence, the model predicts choice based on a mixture of currencies. The reward mixture-of-currencies logistic was the AIC overall best fit (mean AIC = 70.51 ± 1.24; protected exceedance probability = 1) and the best fit in 248 out of 422 participants. The same reward mixture-of-currencies model was the BIC overall best fit (mean BIC = 98.48 ± 1.18; protected exceedance probability = 1) and the best fit in 160 out of 422 participants (see Supplemental Table 1).

**Table 1.**
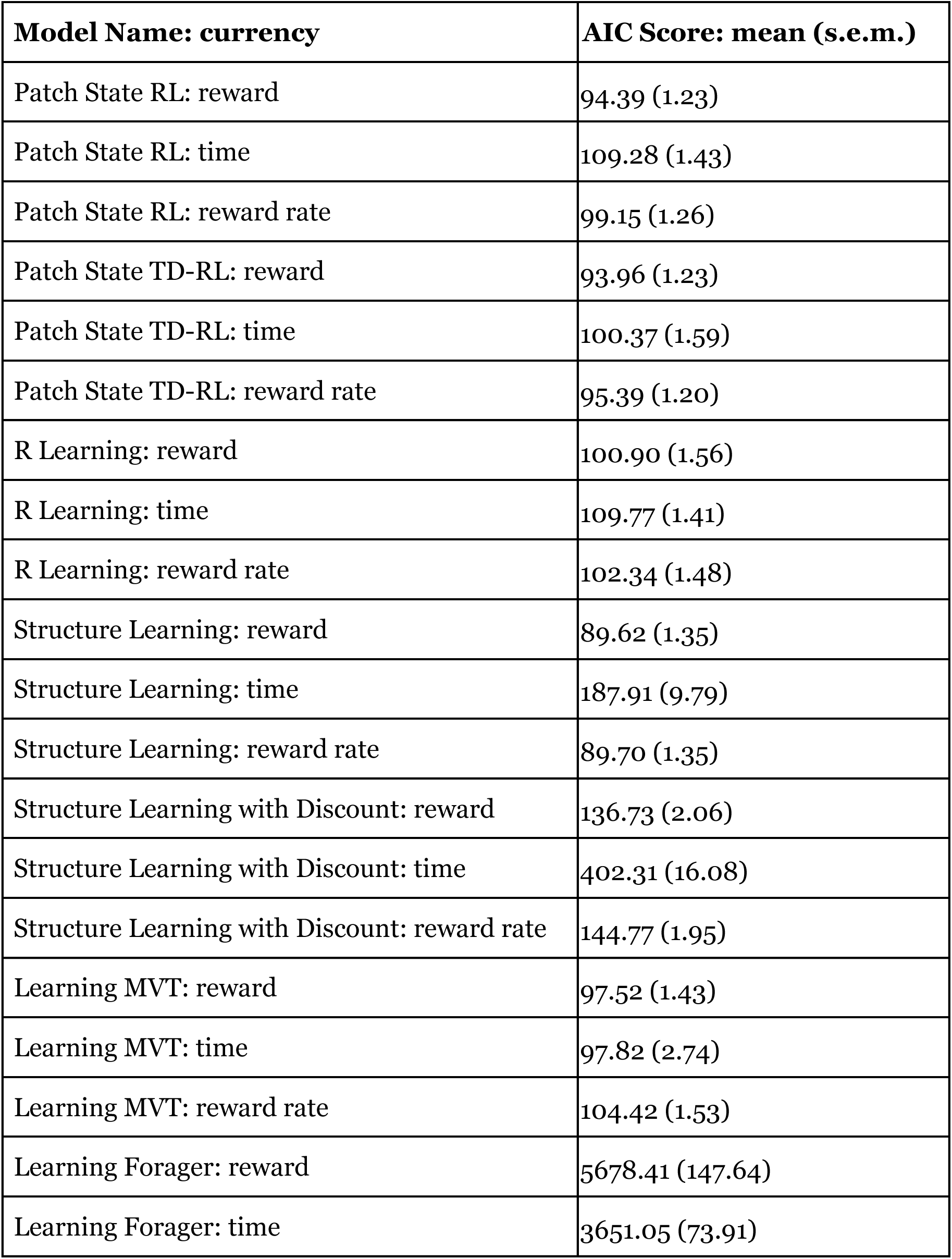
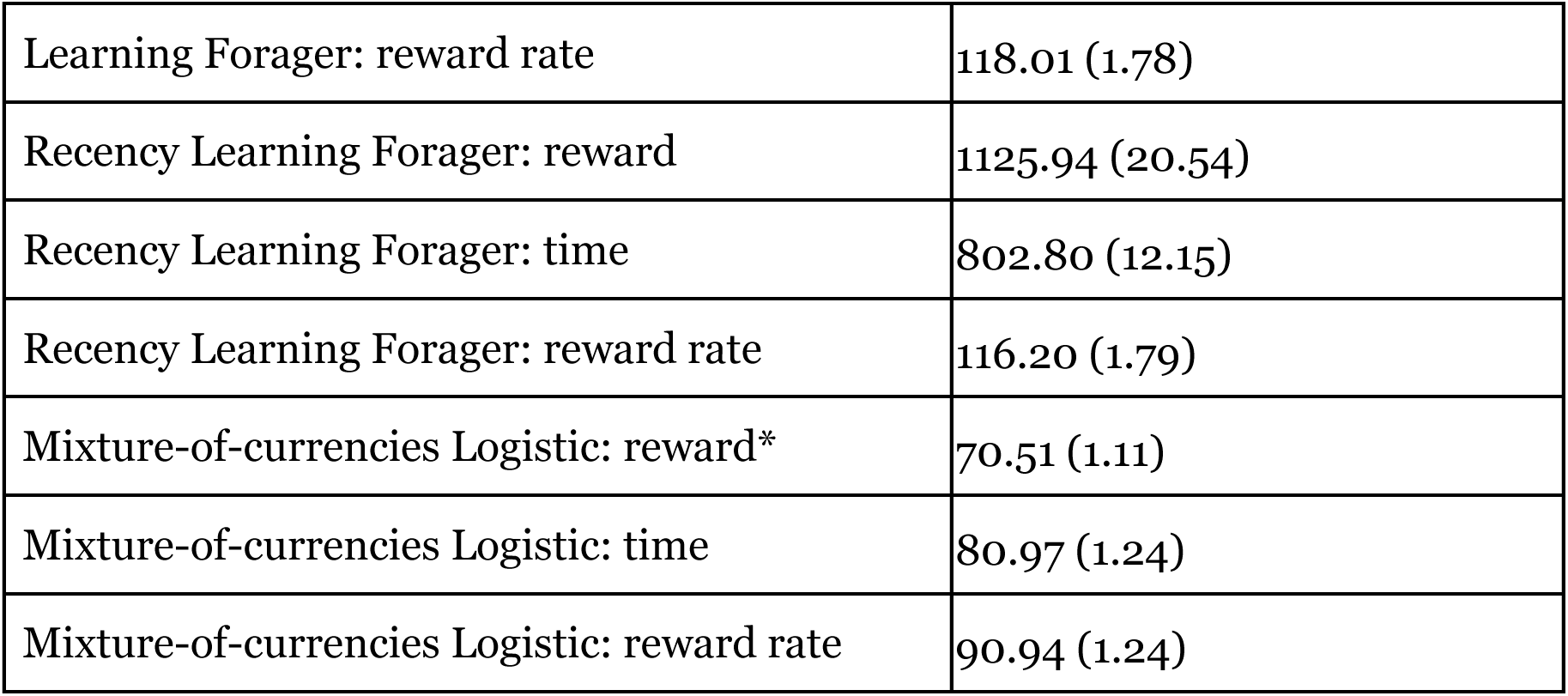
Model fit results for AIC. *: best-fit model overall. See methods for descriptions of each model.

The information term in the best fit model is computed from the decrease in entropy as participants experience the drop in reward from choosing to stay in a patch. To determine if including this term helps explain the observed changes in patch residence time across blocks, we next correlated the best-fit beta coefficient for information with the best-fit OLS slope of patch residence time against patch number in block. We found that the beta coefficient for information for the 1 sec timeout condition correlated with the slope for 1 sec timeout patches (ρ = 0.14, p < 0.005) and the beta coefficient for information for the 5 sec timeout condition correlated with the slope for the 5 sec timeout patches (ρ = 0.32, p < 1×10^-10^). This suggests that one reason the information threshold model performs so well is that it is able to explain changes in patch leaving decision dynamics.

To validate this conjecture, we conducted simulations assessing how well the model fit the observed behavior. We simulated 1000 iterations of the task using each participant’s best fit parameters. These simulations successfully recovered the observed decrease in patch residence times over blocks (Fig. 4). The model accurately captured the temporal dynamics, with longer simulated residence times in earlier patches compared to later ones, providing evidence that the information threshold model explains both overharvesting and its diminution over time. The simulations validate the role of information as a driver of patch foraging behavior.

### Monkey Patch Leaving Task and Behavior

We next turned our modeling lens on monkey foraging behavior to determine if the decision dynamics observed in humans persist in other primates, suggesting generalizability. Monkeys provide an important comparison to humans given their evolutionary proximity and sophisticated foraging capacities (Passingham and Wise 2012). If monkeys display similar dynamics, it would suggest these effects stem from a shared mechanism guiding decisions to depart patches. In addition, testing monkeys enables behavioral data collection after extensive training on the task (>10,000 trials) and assessment of the persistence or attenuation of decision dynamics over time.

We used a previously collected patch foraging dataset (Barack et al. 2017). Two male rhesus macaques (*M. mulatta*) performed a patch leaving task that recapitulated the basic format of the human patch leaving task above (Fig. 5). Monkeys first fixated a central fixation cross for a variable amount of time. Two targets then appeared, a small blue rectangle signaling a ‘stay’ decision and a gray rectangle signaling a ‘leave’ decision. If monkeys chose to stay, they experienced a brief delay followed by a squirt of juice. This was followed by an ITI and the same choice between staying and leaving, except the juice reward for staying decreased by a small amount. If monkeys chose to leave, they experienced a timeout ‘travel time’ delay, signaled by the height of the gray rectangle. Unlike the human experiments, which separated timeouts into blocks, monkeys experienced randomized timeouts between 0.5 to 10.5 sec on each patch. This was followed by an inter-trial interval and a new decision to stay or leave, albeit the two options had switched locations, a new height for the gray rectangle was randomly selected between 0.5 and 10.5 seconds, and the stay reward was reset to its full value.

**Figure 5.**
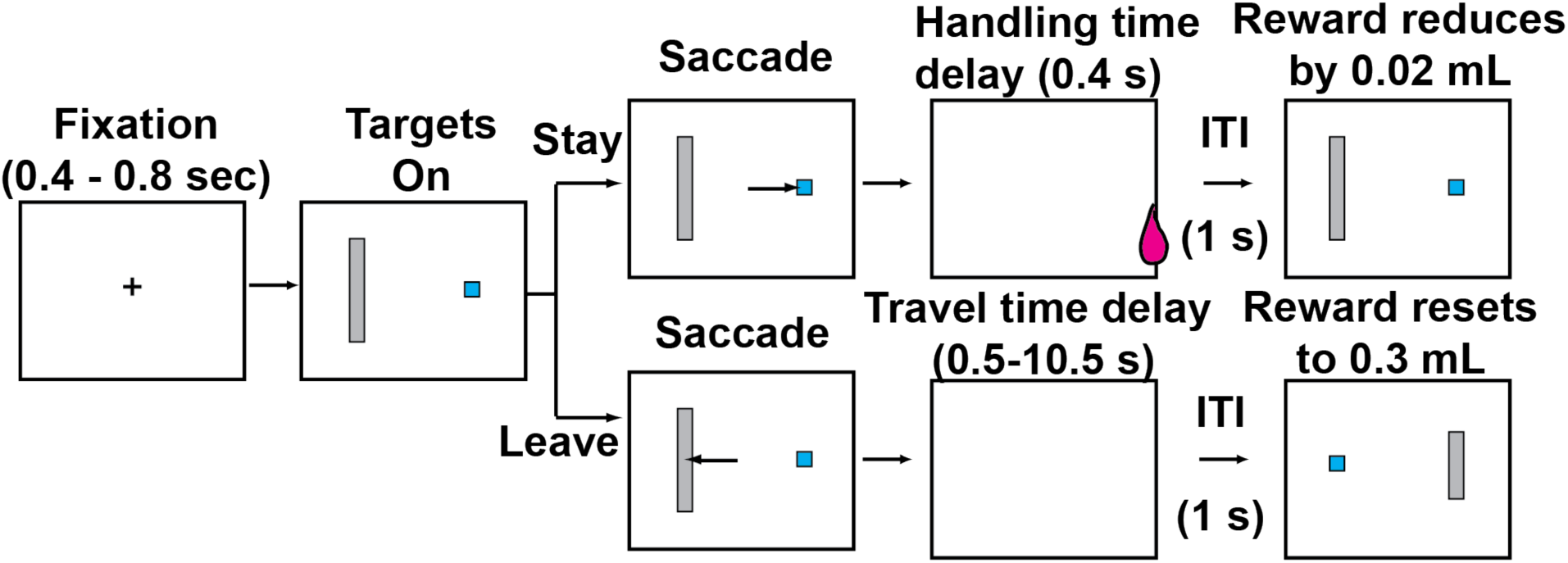
Monkey patch leaving task.

Mirroring human behavior, monkeys stayed longer in patches when the timeout delay was longer (as reported previously in (Barack et al. 2017); OLS; both monkeys, mean β_slope_ = 1.33 ± 0.12 sec in patch / sec delay, Student’s t-test, t(df = 39) = 11.30, p < 1×10^-13^; M1: mean β_slope_ = 1.09 ± 0.19, t(df=19) = 5.60, p < 0.0001; M2: mean β_slope_ = 1.56 ± 0.12, t(df=19) = 13.18, p < 1×10^-10^). Both monkeys also earned more rewards in longer timeout patches (OLS; both monkeys, mean β_slope_ = 17.09 ± 2.23, t(df=39) = 7.65, p < 1×10^-8^; M1: mean β_slope_ = 7.66 ± 1.88, t(df=19) = 4.07, p < 0.0032; M2: mean β_slope_ = 26.52 ± 2.76, t(df=19) = 9.63, p < 1×10^-8^).

Critically, monkeys also exhibited patch residence time dynamics matching the human results – more quickly leaving patches later in sessions compared to earlier. Both monkeys tended to leave patches sooner later in sessions compared to earlier (Figure 6; OLS regression of patch residence time concatenated across days against patch number in session; M1: β_slope_ = -0.14 ± 0.018 sec / patch number in session, p < 1×10^-12^; M2: β_slope_ = -0.036 ± 0.0085 sec / patch number in session, p < 0.0005).

**Figure 6.**
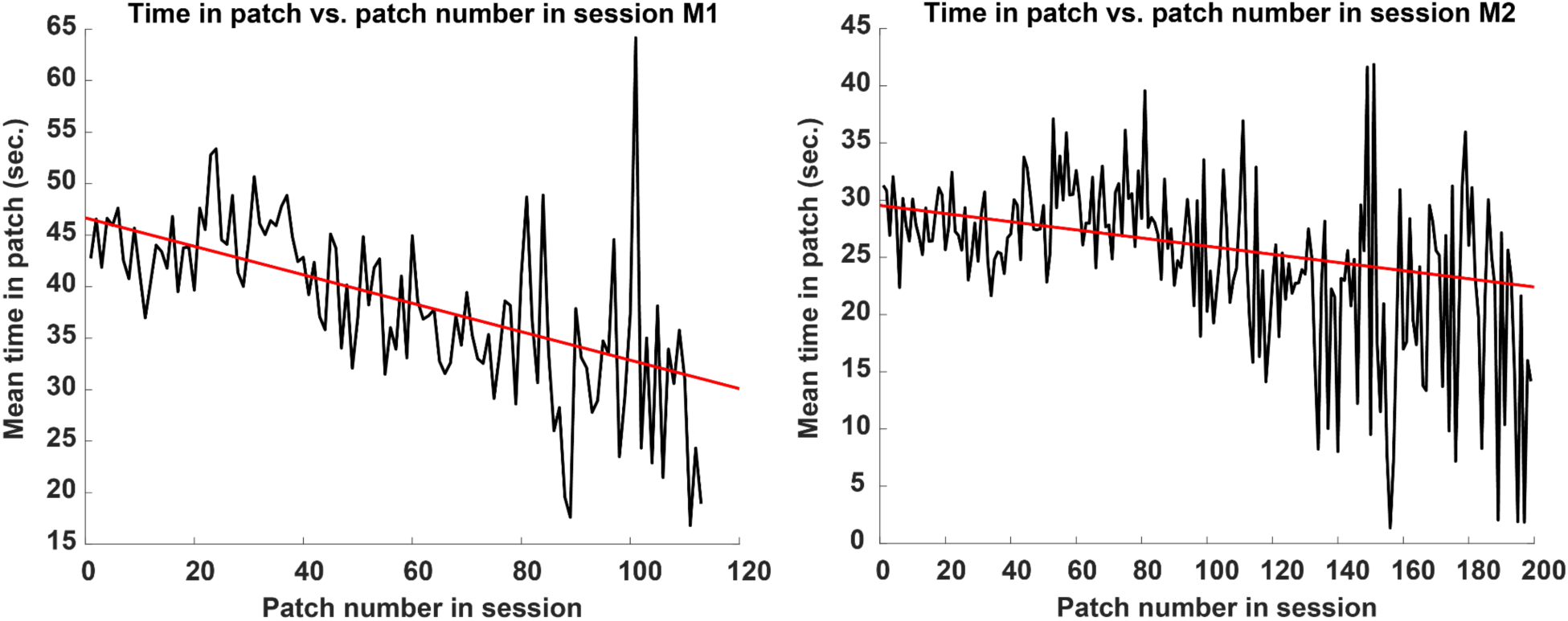
Monkeys spend less time in patches with increasing experience. Mean time in patch in seconds vs. patch number in session. Left: monkey 1; right: monkey 2. Black lines: data; red lines: best-fit OLS linear regression.

Unlike humans, monkeys did not tend to increase their reward rates over the course of a session. One monkey showed no significant tendency to increase or decrease earned reward rates (OLS regression of z-scored patch reward rate concatenated across days against patch number in session; M1: β_slope_ = -0.0010 ± 0.0011 (juice/sec) / patch number in session, p > 0.37), whereas the other monkey decreased earned reward rate (M2: β_slope_ = -0.0036 ± 0.0006 (juice/sec) / patch number in session, p < 1×10^-5^). To further investigate this species difference, we performed two additional analyses. First, we regressed response time on trials when monkeys chose to depart patches against patch number in session. M1 showed no impact on response time (OLS regression of leave trial response time concatenated across days against patch number in session; M1: β_slope_ = - 0.00040 ± 0.0015, p > 0.78), whereas M2 tended to respond more slowly over the course of the session (M2: β_slope_ = 0.00094 ± 0.00026, p < 0.005). Second, we regressed the total duration of leave trials as a fraction of the total time in patch against patch number in session. Leave trial duration includes not only the leave trial response time from the previous analysis, but also all within trial event durations like fixation times as well as any delays from the monkey related to initiating the trial by acquiring the first central fixation. This analysis revealed that for both monkeys leave trials constituted an increasingly greater proportion of total patch time as sessions progressed (OLS regression of leave trial as fraction of total time in patch concatenated across days against patch number in session; M1: β_slope_ = 0.00058 ± 0.00014, p < 0.001; M2: β_slope_ = 0.00037 ± 0.000074, p < 1×10^-5^). By contrast, human participants generally did not exhibit changes in response times on leave trials (106 / 422 participants showed significant changes, p < 0.05) or in the proportion of total time in patch occupied by leave trials (124 / 422 participants significant, p < 0.05). When they did, they tended to respond more quickly (93 / 106 of significant participants with β_slope_ < 0) and leave trials tended to take up increasingly less total time in patch (82 / 124 of significant participants with β_slope_ < 0). In sum, unlike the humans, monkeys tended to take longer to leave a patch and more of the total time in patch was spent on leave trials as sessions wore on, both reducing reward rates. These differences may be due to reward modality (points in the case of humans, squirts of juice for monkeys), reward value (the value of juice goes down as monkeys become satiated, whereas the value of points may not diminish), or task focus (monkeys performed the task until they were satiated, whereas humans completed all patches within 8 minutes).

### Computational Modeling of Decisions Made by Monkeys during Foraging

We next fit the same nine models and three currencies described above to the monkey foraging choices. We report the results for each monkey separately in Table 2. The best-fit model was the mixture-of-currencies logistic that utilized both rewards and information to predict patch leave decisions (M1: mean AIC = 24.53 ± 6.12; M2: mean AIC = 37.59 ± 19.44; protected exceedance probability across sessions = 1 for both monkeys). The model was the best-fit in 20/20 sessions in M1 and 19/20 in M2. These findings were confirmed using BIC (see Supplement). In sum, the best-fit currency and the best-fit model were the same for monkeys and humans.

**Table 2.** Monkey model fit results. . *: best-fit model overall. See methods for descriptions of each model. Each column reports the mean AIC score (s.e.m.) across sessions for that monkey.

| **Model** | **AIC Score: M1** | **AIC Score: M2** |
| --- | --- | --- |
| Patch State RL: reward | 177.83 (15.29) | 278.86 (22.78) |
| Patch State RL: time | 177.80 ( 15.39) | 278.6335 (22.83) |
| Patch State RL: reward rate | 165.51 ( 16.24) | 278.13 ( 22.88) |
| Patch State TD-RL: reward | 151.54 ( 12.96) | 256.45 ( 22.57) |
| Patch State TD-RL: time | 159.77 ( 16.03) | 274.86 ( 21.29) |
| Patch State TD-RL: reward rate | 141.32 (9.37) | 260.88 ( 20.37) |
| R Learning: reward | 166.49 (14.42) | 260.76 (23.56) |
| R Learning: time | 166.29 ( 12.79) | 269.66 ( 21.45) |
| R Learning: reward rate | 157.00 ( 12.13) | 265.00 (23.22) |
| Structure Learning: reward | 163.33 ( 11.18) | 274.42 ( 21.82) |
| Structure Learning: time | 186.10 (15.04) | 282.04 ( 22.60) |
| Structure Learning: reward rate | 167.28 ( 11.90) | 274.88 ( 21.87) |
| Structure Learning with Discount: reward | 318.89 (22.97) | 469.70 (40.30) |
| Structure Learning with Discount: time | 345.65 (28.03) | 479.44 ( 48.43) |
| Structure Learning with Discount: reward rate | 339.57 ( 32.85) | 504.81 ( 38.56) |
| Mixture-of-currencies Logistic: reward* | 24.53 (6.12) | 37.59 ( 19.44) |
| Mixture-of-currencies Logistic: time | 131.79 ( 13.50) | 244.78 ( 19.75) |
| Mixture-of-currencies Logistic: reward rate | 147.24 (12.96) | 268.27 ( 21.41) |
| Learning MVT: reward | 382.27 (30.08) | 574.16 (45.21) |
| Learning MVT: time | 380.31 (30.00) | 572.11 (45.09) |
| Learning MVT: reward rate | 281.53 (17.47) | 518.27 (43.06) |
| Learning Forager: reward | 201.00 (16.25) | 309.41 (24.44) |
| Learning Forager: time | 187.17 (15.09) | 283.09 (22.64) |
| Learning Forager: reward rate | 622.82 (97.72) | 1,330.50 (116.55) |
| Recency Learning Forager: reward | 200.73 (16.57) | 309.09 (24.56) |
| Recency Learning Forager: time | 184.8332 (15.1513) | 280.83 (22.58) |
| Recency Learning Forager: reward rate | 616.31 (96.52) | 1,322.69 (116.49) |

## 3. Discussion

Foraging is a basic competence for all animals that seek resources (Stephens and Krebs 1986) and has been proposed as a potent selective pressure on the evolution of primate intelligence (Passingham and Wise 2012). A core feature of foraging is the decision to continue to exploit depleting resources or to explore for new ones. Recent research has focused on how humans and other primates make these decisions. Here, we sought to answer two key questions: what inputs do primates use to make these decisions and how do they transform those inputs into decisions? To answer these questions, we collected behavior from hundreds of human participants engaged in a virtual foraging task. Consistent with prior studies, humans foraged longer in patches and gathered more rewards when the timeout to replenish a patch was longer, and they over-harvested from patches, staying longer than the optimal time for maximizing rewards harvested. We discovered decision dynamics in patch leave choices, with longer patch residence times earlier in runs compared to later. To explain these findings, we fit 27 different models to observed choice behavior, with nine different proposed transformations operating over three different currencies. We discovered that humans base foraging decisions on obtained rewards. We also discovered that a relatively simple logistic transformation over a mixture of currencies, which included information, best explains leave decisions. To understand the adaptive significance of our findings, we fit the same models to a dataset of monkey foraging behavior. Monkeys’ decisions were also best described using rewards as currency, and monkeys also appeared to use a logistic transformation of a mixture of currencies to make patch leaving decisions.

We considered three canonical foraging currencies: reward, time, and reward rate. From an evolutionary perspective, such as that taken by ecology, a proxy for long-term fitness is the appropriate currency. These three currencies are motivated by considerations of this long term fitness. While maximization of reward rate is often considered the most reasonable proxy for long-term fitness (Stephens and Krebs 1986), the best fit currency for humans was reward. This effect may arise because humans had a short-term focus, seeking to maximize earnings within the fixed duration of the task rather than considering longer-term rates (Heilbronner and Hayden 2013). The best fit currency for monkeys was also reward. Besides a similar short-term focus, this result may be accounted for by reports that animals have trouble tracking time in laboratory tasks (Pearson et al. 2010; Blanchard et al. 2013; Kane et al. 2019). Specifically, non-human animals fail to understand forced post-reward delays, such as inter-trial intervals, which are common in the lab but absent in the wild (Bateson and Kacelnik 1996). The monkeys fail to take into account these delays in their decision calculus, making time and reward rate, which is computed in part from time, worse at explaining their choice behavior, thereby explaining why the currency in the best-fit model was reward.

Our best-fit model included a fourth currency, information. At first pass, this currency does not seem to maximally contribute to long-term fitness. We hypothesize that including this currency allows both humans and monkeys to prepare for the future by learning the structure of the current environment.

The finding that the mixture-of-currencies logistic model best fits the data contributes to a growing understanding of how foraging decisions are made. Changes in foraging behavior have long been attributed to learning, whether optimally (McNamara and Houston 1985) or under constraints such as memory for past foraging success (Ollason 1980). Here, we quantified how monkeys and humans learn the statistics of changes in the environment’s reward structure, which helps explain the dynamics of patch leave decisions. At the beginning of sessions, participants are ignorant of the size or variance of these changes. Continuing to harvest rewards from patches beyond the optimal leave time provides crucial information about the statistics of the environment. By contrast, later in sessions foragers have a better understanding of these changes.

The study of foraging has investigated various explanations for overharvesting. One hypothesis is that overharvesting results from a lack of understanding of post-reward delays (Pearson et al. 2010; Blanchard et al. 2013). By contrast, Kane and colleagues (2019) demonstrate that animals are sensitive to post-reward delays, suggesting that they understand task structure. However, there may be other task parameters that could be learned by sampling from a patch past the point of optimal intake rates. We found that including this information in our models provided a simple succinct explanation of the dynamics of patch harvesting. Indeed, dynamically adjusting thresholds for exploration during foraging is observed in numerous species besides humans (Le Heron et al. 2020; Harhen and Bornstein 2023), including starlings (Cuthill et al. 1990), bumblebees (Hodges 1985), and baleen whales (Piatt and Methven 1992), amongst others. Besides competition, nutrient availability, or type of food, these dynamic adjustments might be rationally explained as information-gathering to improve estimates of the availability of resources or features of the environment (Pyke et al. 1977; McNamara and Houston 1985). Two other explanations for overharvesting are structure learning and influences of recent choices and outcomes in serial decision making. Harhen and colleagues (Harhen and Bornstein 2023) discovered that overharvesting in environments with more than one type of reward function can be explained by structural inference of different states of the environment. There are two drawbacks to this account. First, overharvesting occurs even in simple environments that feature a single patch value decrease function, as reported here and in numerous other papers (Hayden et al. 2011; Barack et al. 2017; Kendall and Wikenheiser 2022). Second, their model may be more complex than needed to explain overharvesting, which may occur due to simple computations over a mixture of currencies, as we found. Static decision models (such as (Lau and Glimcher 2005)) assume serial choice effects are due to either choice or reward history. However, such serial effects may be dynamic, with the impacts of choice and reward history varying over time. In order to explain dynamic decision effects, a serial decision model would need to treat the effect of, say, a choice or reward two trials before the current trial separately at the beginning compared to the end of runs. This is tantamount to the trivial, over- parameterized model where each choice and outcome receives its own best-fit coefficient.

In contrast to these explanations, non-reward priorities such as information (Bromberg- Martin and Monosov 2020) may shape decision dynamics of staying or leaving patches. Here, we reported that accumulating information about the structure of the environment can give rise to dynamic decision effects on explore-exploit tasks. Because information naturally decreases over the duration of the task, the acquisition of information can readily explain the dynamics of overharvesting, including what appear to be the effects of recent choice or reward history.

Our best-fit model used a mixture of currencies–reward and information–but a simple logistic computation to generate choices. Decision dynamics generally can be explained in two ways, corresponding to computation and currency. Computation explanations posit increasingly sophisticated transformations of a currency in order to produce decision dynamics like the change in overharvesting we observed over the course of sessions. Currency explanations posit that decision computations use multiple different variables to produce choices. In short, there is a computation-currency tradeoff in explanations of foraging. Decision dynamics may result from increasingly complex computations over some currency or from changes or expansions of currencies. While recent explanations of these effects in foraging have focused on increasing the complexity of the computation (Constantino and Daw 2015; Harhen and Bornstein 2023), we demonstrate that an alternative and perhaps better explanation of the dynamics of foraging decisions utilizes multiple currencies.

Our findings should be approached with some caution. Neither task involved any great metabolic costs and no time costs at all outside the delays; hence, we could not address debates over whether reward rates or efficiency (rewards gained compared to rewards lost), or whether gross or net reward, are the relevant measures (Kacelnik and Houston 1984; Schmid-Hempel et al. 1985; Ydenberg et al. 1994; Bateson and Kacelnik 1996). Further, as noted, our findings are at odds with the widely accepted view that reward rates are the relevant currency for foraging. More careful manipulation of time delays during the task (such as found in (Blanchard et al. 2013) or (Kane et al. 2019)) or shifting to a freely-moving foraging environment might provide insights into this discrepancy. Finally, humans improved their reward rates over sessions whereas monkeys did not. This appeared to be due to differences in how the two species behaved on leave trials. Monkeys were less eager to leave patches as sessions wore on, taking longer to choose to depart, and leave trials occupied an increasingly greater fraction of total time in patch. These differences may be due to interspecific differences in task focus or the effects of satiety (Grether et al. 1992).

Our study lays groundwork for understanding how humans and animals gather information even when making simple choices. Skilled information seekers like primates in particular may always be dynamically adjusting their models of the world to improve long-run payoffs. This adjustment process helps illuminate why humans and other animals make exploratory decisions that seem suboptimal yet ultimately produce adaptive behavior.

## 4. Conclusion

Computational modeling revealed that a simple threshold model using both reward and information as currencies best described foraging decisions. On this computational model, participants decide to stay or leave a patch based on a combination of the recent rewards as well as the amount of information they anticipate gaining from the outcome of their choices. As participants forage for rewards, they gather decreasing amounts of information as the environment becomes better learned, helping explain overharvesting as well as the dynamics of patch residence times.

## 5. Methods

### Task

We recruited participants (n = 506) via Prolific to complete a virtual patchy foraging task programmed in Javascript followed by a number of surveys (results reported elsewhere). After preprocessing, 42 individuals were excluded from final analyses for the following criteria. First, they either failed pre-defined survey attention checks or did not complete every survey. Further, participants were excluded if they completed less than 25 trials (i.e., an exploit or explore decision) throughout the session. We also excluded participants that were believed to have misunderstood the instructions given their comments (e.g., “I don’t know if I understood it” or “I did not understand until the last round”). Finally, we excluded an additional 39 individuals who failed to complete at least 3 patches in both short (1 sec) and long (5 sec) timeout conditions. We enforced this last constraint to ensure participants were engaged with the task and completed enough patches to assess whether patch residence times changed over the course of blocks. Of the final 422 participants (µ_age_ = 44.66 y.o., σ_age_ = 15.98), 210 identified as “male”, 207 identified as “female”, and 5 identified as “other”.

In the task, a “valid patch” begins with an exploit decision, in which the user collects berries, and ends with an explore decision, in which the user travels to the following patch. Note that any patches where the participant immediately explored was considered a valid decision, too, and thus included in the analysis. Any “incomplete patch” that did not end in the user leaving the patch (explore) was removed. These incomplete patches occurred at the end of session blocks.

Participants were sent to the training and task via a link in Qualtrics after completing an informed consent form and several questionnaires. Before starting, participants received detailed written instructions on the task and watched a video demonstration. To familiarize themselves with the task, they also completed a 1 min training (30 sec per block type) before starting the actual experiment. The instructions prior to the training were (emphasis in bold and color as in original):

> “In this experiment, your goal is to **collect as many berries as you can** in the amount of time you are given.

> First, hover over the center box until it fills up **completely**. You may have to **wiggle your cursor** once for the center box to recognize your mouse.

> Then, on each trial, you will either choose to **stay** at the bush you are at or **leave** the current bush and travel to a fresh bush.

> To **pick berries** from the current bush, hover your cursor over the bush next to the center box.

> The number of berries you pick at each attempt will be indicated next to the bush. As you keep picking from the same bush repeatedly, the number of berries you pick at each attempt will go down as you will be depleting this bush.

> To **travel** to the next bush, mouse over the box on the opposite side of the bush. You can choose to leave the current bush to travel to a fresh bush at any time. When you do that, you will need to wait while you travel to the next bush.

> However, once you arrive at the new bush, the amount of berries you pick at each harvest will reset to the initial value.

> You will play different blocks in this game. The environment will change between these blocks.

> There is an **unlimited number of bushes** in the environment, but a **finite amount of time**. Try to collect as many berries as you can!”After completing the experiment, participants returned to Qualtrics to finish further questionnaires. Data were stored on a secure AWS server.

During the task, participants collected as many berries as possible in 8 min (4 blocks, 2 min each). The reward function for the task started at 7 plus Gaussian noise for both long and short timeout patches:

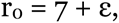

where ε ∼ N(0, 0.25) and then, for each choice to harvest reward, decremented by 0.5 plus noise:

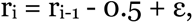

where again ε ∼ N(0, 0.25). At the end of the experiment, participants received 0.3226 cent per berry collected (up to $3 in total) in addition to their base participation compensation of $4.

Participants were randomly assigned to one of two orders of timeouts by block: order 1 [block 1: 1 sec timeout, block 2: 5 sec, block 3: 1 sec, block 4: 5 sec] or order 2 [block 1: 5 sec timeout, block 2: 1 sec, block 3: 5 sec, block 4: 1 sec].

### Monkey Task, Training, and Care

Two mature (aged ∼6-9 years) male rhesus macaques (*M. mulatta*) participated in the foraging experiment. Monkeys were single housed in cages in a colony room with other monkeys. Monkeys received daily enrichment and biannual health check-ups. After initial behavioral training, a head-restraint prosthesis (titanium; Crist Instruments) was implanted using standard aseptic surgical techniques. All surgeries were performed in accordance with protocols approved by the Duke University institutional animal care and use committee and were in accord with the Public Health Service *Guide to the Care and Use of Laboratory Animals*. Monkeys were anesthetized using isoflourane, received analgesics and antibiotics after surgery, and permitted a month to heal before any further experiments.

During training, monkeys’ access to fluid was controlled outside of experimental sessions. Custom software written in MATLAB (Mathworks, Natick, MA, USA) using Psychtoolbox (Brainard 1997) controlled stimulus presentation, reward delivery, and recorded all task and behavioral events. Horizontal and vertical eye traces were sampled at 1000 Hz by an infrared eye-monitoring camera system (SR Research, Osgoode, ON) and recorded using the Eyelink toolbox (Cornelissen et al. 2002). Solenoid valves controlled juice delivery. All data were analyzed using custom software written in MATLAB.

Our task simulates a patch-leaving problem by presenting the animal with a two-alternative forced choice decision between continuing to forage at a depleting resource and waiting to replenish the resource. To begin the trial, the animal fixated (± 0.5°) on a centrally presented cross for a random fixation time drawn from a uniform distribution (400 – 800 ms). If the animal prematurely shifted his gaze from the fixation cross before exhausting this time, the fixation clock resets to zero. If the animal exhausted the fixation time, the fixation cross was extinguished and the targets appeared, a small blue rectangle and a large gray rectangle, one each on the left and right side of the screen. The animal could make a choice by aligning gaze with a target and holding it there for 250 ms.

If the monkey selected the blue rectangle, he was permitted to freely look about while the rectangle shrank at 65 pixels/s until it disappeared. This shrink time simulated the ‘handling time’ for the food item, and was constant across all trials and reward sizes. At the end of this handling time period, the animal received a squirt of juice, followed by a 1 second intertrial interval (ITI) and the reappearance of the fixation cross. The reward size for the first trial in patch was always ∼0.30 mL of juice. As the animal continued to select the blue rectangle (‘stay in patch’ decision), the amount of juice associated with that choice dropped each trial by ∼0.02 mL of juice. After a series of stay in patch decisions, the animal typically decided to select the gray rectangle (‘leave patch’ decision). After selecting that option, the monkey was free to look about while the gray rectangle shrank also at 65 pixels/s. The height of the gray rectangle signaled the time-out penalty for leaving the patch and did not vary so long as the animal continued to stay in the patch. Once the monkey chose to leave, the gray bar shrank, which was followed by a 1 second ITI and the reappearance of the fixation cross; no juice was delivered for this choice. On the first trial in the ‘new patch’, the juice reward associated with the blue rectangle was reset to its full amount, the height of the gray bar was selected randomly from the distribution of 0.5 – 10.5 s, and the locations of the targets were switched. Each session was limited to one hour.

### Data Analysis

All behavioral data from both species were analyzed using custom software in MATLAB. For patch residence time comparisons, we averaged the median patch residence time across participants, in order to control for outliers. We used within- participant paired t-tests to compare the average patch residence time for 1 sec and 5 sec timeout patches. Total time in patch was determined by the amount of time from the first choice to collect rewards in a patch to the decision to depart a patch.

For reward-related comparisons, we used both cumulative reward (which has in the past been the computationally relevant variable for decisions to harvest rewards in patch; (Barack et al. 2017) and reward rate (the classic currency for foraging choices in optimal foraging theory; (Stephens and Krebs 1986). Cumulative reward was defined as the running sum of berries (humans) or the total msec of open solenoid (monkeys) collected within a patch, and reward rate was defined by dividing cumulative reward by the cumulative time in patch. We used within-participant paired t-tests between timeouts to compare cumulative rewards and reward rates.

To compare human to monkey behavior, response times on leave trials and leave trial durations were both analyzed. Response times were defined generally as the time from the end of the ITI box filling to time of choice, when participants moved their mouse over either the stay or leave option. These response times were computed for each patch and then regressed against patch number in session using OLS. Leave trial durations were defined as the time from ITI box onset to time of choice. Leave trial durations were divided by total time in patch and then regressed against patch number in session using OLS.

For the monkeys, time in patch was determined by the amount of time from the first choice to collect rewards in a patch to the decision to depart a patch. For the regression of time in patch vs. patch number in session, time in patch for each patch was concatenated across sessions and regressed against the patch number in session.

Cumulative reward was defined as the running sum of the juice reward time collected within a patch, and reward rate was defined by dividing cumulative reward by the cumulative time in patch. For the regression of reward rate vs. patch number in session, reward rates were first z-scored by session, and then reward rates by patch concatenated across sessions were regressed against the patch number in session.

Leave trial response times and durations were both analyzed. Response times generally were defined as time from end of fixation (i.e., go cue) to time of choice (i.e., registration of selecting either the stay in patch or leave patch box), including for leave trials. These were computed for each leave decision, concatenated across sessions, and then regressed against patch number in session using OLS. Leave trial durations were defined as the time from onset of fixation to time of choice. These durations were divided by the total time in patch, concatenated across sessions, and then regressed against patch number in session using OLS.

### Optimal Behavior

First, an exponential curve was fit to the cumulative intake curve defined by the gain function (*fit* function in MATLAB). The gain function g(t) for time in patch t is well- described by an exponential curve, which can be used to solve the optimal leave time from first principles (cf. Cowie 1977):

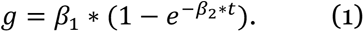

While this continuous and differentiable function is at odds with the discrete series of actual rewards, using an exponential permits a first principles solution to the optimal patch stay time. To find the optimal leave times, the first derivative of g(t) is set equal to the gain function divided by the average timeout and time in patch:

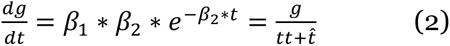

for timeout tt and optimal leave time. Substituting (1) in for g and rearranging terms yields

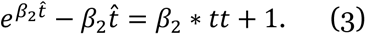

We next used the McLaurin expansion for e, eliminated constants, dropped the fourth order and higher terms, subtracted out the first order term, and rearranged to arrive at

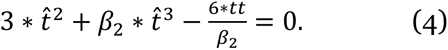

We used the best-fit β_2_ from our exponential fit (1) to the series of average returns. Then, using the short (tt = 1 sec) and long (tt = 5 sec) timeouts in turn, we determined the optimal leave time by solving (4) with numerical methods (*fmincon* in MATLAB subject to nonlinear constraints determined by the McLaurin expansion).

### Model Fitting

We fit a number of models inspired by different formal frameworks to decisions to stay or leave a patch. A large number of models (n = 27) were fit using *fmincon* in MATLAB, with parameter estimates for each participant obtained by using standard maximum likelihood estimation. All models were assessed using both AIC and BIC to confirm best-fits, and all models included a catch probability for leaving patches on the first trial in a new patch as well as a side bias term to capture the influence of the location of the options on screen. Finally, protected exceedance probabilities were computed from AIC and BIC scores using custom MATLAB software adapted from the *SPM* function *spm_BMS* (Stephan et al. 2009; Rigoux et al. 2014).

#### Model fit score

After finding the maximum likelihood estimates (MLE) for the best-fit parameters, defined as the maximum of the log of the likelihood scores

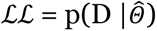

for log likelihood score ℒℒ, data D and parameter estimates *Θ̂*, model fit scores were computed using AIC

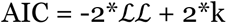

for number of parameters in the model *k*. Model fit scores were also computed using BIC

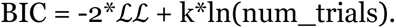

These scores are reported in Table 1 and Supplemental Table 1.

#### Currencies

We fit our models using three different currencies, variables that are input into the decision computations, one of the nine models discussed below, used to predict choices. The three currencies are reward, time, and reward rate. Reward was defined as the reward received on the previous trial, points in the case of humans and msec of open solenoid for juice delivery in the case of the monkeys. Time was defined as the total time in patch: time from trial onset (onset of either the ITI box for humans or the initial fixation cue for the monkeys) until the time of the decision (end of the ITI box filling for humans or disappearance of the fixation cue for the monkeys). While it is perhaps unusual to use time as a currency, timing rules for patch departure are not uncommon (see, e.g. Constantino and Daw 2015). Reward rate was defined as the cumulative sum of the rewards in the current patch divided by the cumulative time in the current patch. See below for details about the information currency used in the best-fitting mixture-of- currencies model.

#### Reinforcement learning models

Reinforcement learning models are inspired by classic Rescorla-Wagner reinforcement learning (Rescorla and Wagner 1972) and draw on computational reinforcement learning (Sutton and Barto 1998). Computational reinforcement learning is a set of techniques for finding optimal policies, rules for making actions in particular states. These models propose that foragers keep a cached value of the leave and stay options and on each trial use that cached value to make a decision to stay or leave a patch. The cached values of the options are updated after foragers receive outcomes from each decision. Participants are assumed to learn these values of staying or leaving a patch over time. Decisions are made by maximizing the long-run return from taking actions in particular states. We fit three reinforcement learning models to participants’ choices.

##### 1. Patch state Q-learner

A Q-learning model for patch foraging. In this model, each choice in a patch has it’s own state-action (or ‘Q’) value associated with it, designated *Q(S = s_i_ | A = a_i_)* for state variable *S* set to the *i*^th^ state *s_i_* and action variable *A* set to the *i*^th^ action. Following Constantino and Daw (2015), we simplified the model such that there was a single value associated with leaving across all states, *Q_leave_*, and a separate value for staying for each n^th^ choice in patch, *Q_stay1_*, *Q_stay2_*, etc. There were separate Q-values for both 1 sec and 5 sec timeout patches, but a single value for leaving across all states and patch types. All Q-values were initialized at 3.5, half the mean of the first reward in patch, to hasten learning.

Patch stay and leave decisions were modeled using a softmax equation for the probability of staying

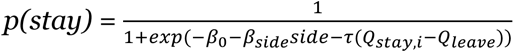

that contained Q-value terms for staying on the *i*^th^ choice in patch, *Q_stay,i_*, the value of leaving *Q_leave_*, a term *side* for the side of the stay choice to capture side bias, and an intercept term. Probability of leaving *p(leave)* was

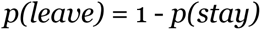

The value input into the softmax was the difference in values, *Q_stay,i_* - *Q_leave_* and were weighted by a choice temperature τ.

We used a standard discounted α-δ update rule (Sutton and Barto 1998) for learning the values of both stay Q-values and the leave Q-value. The value of leaving a patch on choice *t* was updated as

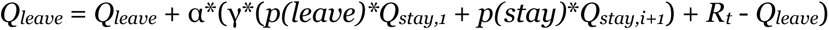

for learning rate α, discount factor γ and reward on the current trial *R_t_*. Because participants earned no rewards on leave trials, *R_t_* = 0 for those trials. The value of staying in a patch on choice *t* for the *i*^th^ choice number in a patch was updated as

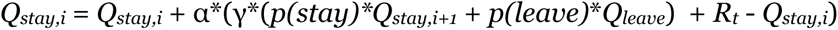

The reward *R_t_* were the points received on that choice. Each patch type updated a separate set of *Q_stay_* values. We used a single learning rate α and a single discount factor γ across both patch types. Parameters α, γ, and β_C_ were ∈ [0 1], whereas the other parameters were allowed to range widely ∈ [-500 500].

The patch state Q-learner, then, had six β parameters: the learning rate α; the discount rate γ; the softmax intercept β_0_; a side bias weight β_S_; a softmax temperature τ; and a catch probability β_1_ for the first trials in patch.

##### 2. Temporal difference(λ) patch state Q-learner

We also fit a version of a temporal-difference learning model with eligibility traces (or TD(λ)) for patchy foraging, after Constantino and Daw (2015). We constructed a backward-looking version of TD(λ). All Q-values were initialized at 3.5, half the mean of the first reward in patch, to hasten learning. In our TD(λ) model, we used the same Q-value update equation for leave decisions as for the patch state Q-learner.

The update equation for the stay decisions, however, was more complex. After making a stay decision and receiving a reward, Q-values were updated starting with the Q-value for the current state *Q_stay,i_* and stepping back through each state’s Q-value to the first state in patch *Q_stay,1_*. The value of staying in a patch on choice *t* for the *i*^th^ state in a patch was updated as

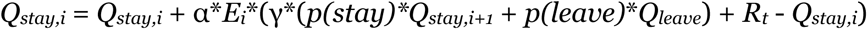

for eligibility trace *E* and other variables are defined above. The eligibility trace *E* was defined as 1 for the current state and

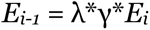

for eligibility parameter λ, discount parameter γ, and patch states *i-1* and *i*. If λ = 0, this simplifies to the patch state Q-learner above. The eligibility trace effectively exponentially discounts the current return and backs up that discounted value to all previous states. The patch stay probability *p(stay)* and patch leave probability *p(leave)* were computed using the same softmax equation above with the same parameters. The TD(λ) Q-learner, then, has the same parameters as the patch state Q-learner plus an additional eligibility parameter λ.

The TD(λ) patch state Q-learner, then, had seven β parameters: the decision noise τ; decision intercept β_0_; side bias β_s_; first trial leave catch probability β_1_; the Q-value learning rate α; the discount parameter γ; and the eligibility trace parameter λ.

##### 3. R-Learner

We fit a third reinforcement learning model using the R-learning algorithm (Schwartz 1993). Like the other reinforcement learning models, the R-learner caches the value of each choice of staying in patch, separately for 1 sec and 5 sec timeouts, and single, separate Q-value for leaving. All Q-values were initialized at 3.5, half the mean of the first reward in patch, again to hasten learning. The R-learning algorithm optimizes the average intake per time step. The model selects choices that maximize the undiscounted long-term return from choices by using the difference between the average return rate and the current outcome in the prediction error δ update instead of simply the current outcome. The algorithm updates the estimate of the average return after each outcome as well, using a second learning rate α_2_.

The R-learning model used distinct update equations for staying and leaving. The update equation for the value of leaving was

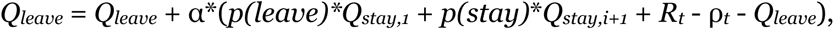

and for the value of staying was

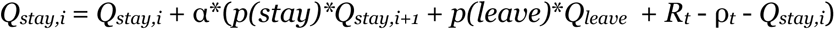

for variables as defined above and average return ρ*_t_* on patches of type *t* (that is, either 1 sec timeout or 5 sec timeout). There was a second update equation for the average return on leaves

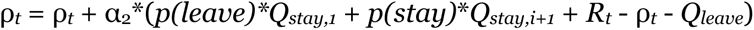

and on stays

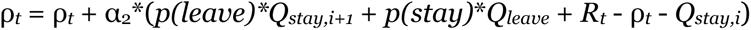

for variable as defined previously. These update equations update the value of the average return by a weighted prediction error across all states, and effectively estimate the average Q-value across all states. The initial values of each ρ were set to 0.

The patch stay probability *p(stay)* and patch leave probability *p(leave)* were computed using the same softmax equation above with the same parameters. The R-learner, then, has the same parameters as the patch state Q-learner, less a discount parameter plus a second learning rate parameter for updating the estimates of the average return.

The R-learner model, then, had six β parameters: the decision noise τ; decision intercept β_0_; side bias β_s_; first trial leave catch probability β_1_; the Q-value learning rate α; and the average return learning rate α_2_.

#### Structure Learning Models

Building on a number of recent models that have investigated how structure in the environment is learned (Gershman and Niv 2010; Collins and Frank 2013), structure learning models model foraging as driven by the search for rewards as well as learning about the environment (Harhen and Bornstein 2023). Even though there was a single reward decrease function in our task, unlike (Harhen and Bornstein 2023), participants may still be inferring the presence of a number of different types of patches due to noise in that function. Here, we adapted and implemented the patch foraging structure learning model from (Harhen and Bornstein 2023).

##### 1. Undiscounted Structure Learner

On our adaptation of their model, the forager infers both the current patch type and the number of patch types using Bayesian inference. Each patch belongs to some patch type, defined by a unique patch value decrease function drawn from a normal distribution

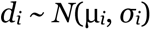

where *i* is the patch type. The observed decreases in the returns from foraging are generated by sampling the distribution of patches *P*(*k*). The initial mean patch depletion rate µ_0_ was set to 0.5 and the initial standard deviation on patch depletion σ_0_ was set to 0.5 as well. On the first trial, no reward drops have been observed, and so the first outcome is assigned to the first patch type *i* = 1, the return *R_t_* was set to an initial value of 7 (matching the average initial reward in patch), the total returns *R_total_* = 0 and total times *T_total_* = 0 as well. Across all choices, patch depletion *d*(*k*) was set to the mean *M*(*k*) for the current patch’s patch type *k*.

Choice probabilities were once again computed using a softmax,

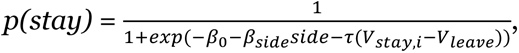

and probability of leaving *p(leave)* was

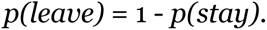

However, the stay values *V_stay,i_* for choice *i* and leave values *V_leave_* were computed differently from the reinforcement learning models. The value of *V_stay,i_* depended on the currency: for reward, *V_stay,i_* = previous trial reward - *d*(*k*); for time, *V_stay,i_* = handling time, the total time to including any ITI, trial initiation time, and median response time to obtain a reward; for reward rate, *V_stay,i_* = (previous trial reward - *d*(*k*)) / handling time, the expected reward rate for the upcoming choice. The value of *V_leave_* also depended on the currency: for reward, the session reward rate was multiplied by the handling time to yield an expected reward; for time, the average time in patch across patches; for reward rate, the session reward rate, defined as the total session reward divided by the total session time. Session rewards *R_total_* were updated after each choice by adding the last trial’s reward and session times *T_total_* were updated after each choice by adding the last trial’s total time. The intercept, side bias, side variables, and temperature parameters of the softmax were defined the same as the reinforcement learning models.

A crucial element of the structure learning model is the update of the distribution of estimated types of patches *P*(*k*). Since this estimate was based on the drops in reward, this distribution was updated only after decisions to stay. Further, for the σ*_k_* update to be valid, participants had to experience at least two drops in reward in a patch, and so *P*(*k*) was only updated after participants experienced two such drops. To update, first, the set of experienced drops D for the current patch was updated by adding the most recent reward decrement. Second, to determine the posterior probability of patch type *k* given the set of drops D, *p*(*k*|*D*), the prior patch type probabilities were computed as the patch count for each type divided by the sum of the patch counts over old types

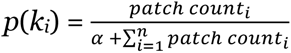

for *n* types of patches, clustering parameter α that partly determines when a new patch is introduced, and the counts for the *i*^th^ patch type *patch_count*_i_. In addition, the probability of a new patch type was computed as

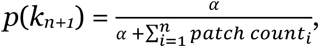

and then *p*(*k*) was normalized to insure it sums to unity. Next, the likelihoods *p*(*D*|*k*) were computed using *normpdf* in MATLAB using the computed patch drop mean µ*_i_* ∈ *M* and variances σ*_i_* ∈ *S* for each patch type *i*. The likelihood of *D* coming from a new patch type was computed using µ_0_ and σ_0_ above. The posterior probability of *D* coming from patch type *i* was then computed as the element-wise product of the priors times the likelihoods, divided by the sum of those products, to normalize.

The set of decreases *D* were assigned simply, selecting the patch type that maximized the probability of *D*. If the final patch in the posterior maximized the probability of *D*, then a new bin was added to the end of *patch_count* with a value of 1, and a new bin added to each vector of patch means *M* and variances *S* set to µ_0_ and σ_0_ respectively from above. If a different patch *k* in the posterior maximized the probability of *D*, then the mean and variance of patch type *k* was updated using standard equations for updating a normal distribution with unknown mean and variance (see Eq. 12 of (Harhen and Bornstein 2023). Because the structure learner model inferred patch types based on the reward functions, we did not fit separate sets of parameters for 1 sec and 5 sec timeout blocks.

In short, each time participants experienced a decrease, the probability of experiencing the set of decreases for the current patch was recalculated using the vectors *M* and *S*. The decreases were assigned to the patch that maximized that probability and the vectors *M* and *S* updated. This in turn set the expected values for the stay *V_stay_* and leave *V_leave_* choices.

The structure learner model, then, had five β parameters: the decision noise τ; decision intercept β_0_; side bias β_s_; first trial leave catch probability β_1_; and the clustering parameter α.

##### 2. Discounted Structure Learner

In addition to the undiscounted structure learner above, we also fit a discounted structure learner, also adapted from (Harhen and Bornstein 2023). The discount structure learner differed from the structure learner in the addition of a discount factor that weighted the value of the leave options, to capture the decreased value of options that are further in the future. The value of *V_leave_* depended on the currency and the discount γ: for reward, the session reward rate was multiplied by the handling time to yield an expected reward and weighted by γ; for time, the average time in patch across patches, also weighted by γ; and for reward rate, the session reward rate, defined as the total session reward divided by the total session time, multiplied by γ.

The discount factor γ was computed after (Harhen and Bornstein 2023). We took the distribution of patch types *p*(*k*) and generated 100 samples from the distribution

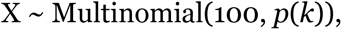

dividing X by its sum to normalize. This sample was used to define the entropy *H_k_* of the patch type distribution

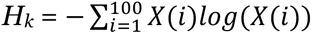

and then used to compute γ

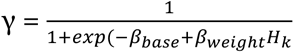

for baseline parameter β_base_ and weight parameter β_weight_. This γ factor in turn discounted the values of the leave options for all three currencies.

The discounted structure learner model, then, had seven β parameters: the decision noise τ; decision intercept β_0_; side bias β_s_; first trial leave catch probability β_1_; the clustering parameter α; and the two γ factor parameters, baseline parameter β_base_ and weight parameter β_weight_.

#### Mixture-of-currencies Model

The mixture-of-currencies model describes participants’ choices as a function of one of the three standard foraging currencies described above (reward, time, reward rate), information, and the interaction between the foraging currency and information. For the humans, blocks of patches were concatenated by timeout condition and modeled separately, as with the other models. This model uses a simple logistic to compute the probability of choosing to stay or leave a patch.

Information for this model was calculated from the decrement in rewards from deciding to stay in patch as follows. Reward decrements Δr were computed by subtracting the reward on trial *t* from the reward on trial *t-1*. Because we wished to use information as a possible motivator for choices, we constructed an expected information variable. Information was computed as the expected change in entropy assuming a normal distribution of reward decrements. The entropy of the normal distribution is

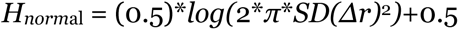

for the standard deviation *SD* of the distribution. We made several accommodations for the first, second, and third trial of patches, when there are too few samples to use this equation, before participants had experienced multiple drops in reward. As described above, the first trial in a patch was fit using a special catch probability across all the models. Before the second choice in a concatenated block of patches, a decrement still has not been experienced. We set the entropy before that choice *H_before_* to

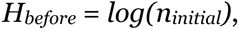

the logarithm of a uniform distribution ranging from 0 to *n_inintial_*, the upper bound on the central tendency of the distribution of reward differences. We set *n_initial_* to 200 to reflect maximal ignorance on the part of the participants regarding the changes in reward. The entropy after the second choice *H_after_* was calculated from

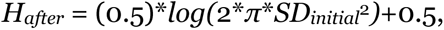

where *SD_initial_* was set to 5, selected as a suitably broad standard deviation to represent participants’ ignorance. For the third choice, only a single reward decrease had been experienced, and so *H_normal_* evaluates either to 0 (biased case) or to *-Inf* (unbiased case), preventing use of the *H_normal_* equation. Instead, for *H_before_*, we once again set the *SD* to 5, yielding

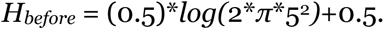

For the entropy after, participants might use their experience of a single reward decrement to estimate the information gained from experiencing a second reward decrease. We used an approximation of the entropy of the normal for a single sample (Ahmed and Gokhale 1989)

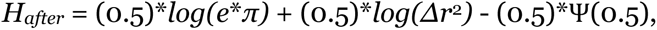

for natural base *e*, single reward decrement *Δr*, and digamma function Ѱ. After the second reward decrement, the entropy before a choice was computed using the equation for the normal distribution above. The entropy after a choice of course is not known until the participant experiences the next reward decrement. To model the expected standard deviation, we made the simple assumption that the expected next decrement was the mean of the previously experienced decrements, 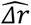, with entropy *H_after_* computed as

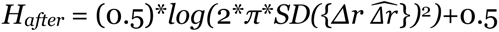

where *H_after_* is a function of the standard deviation *SD* of the union of the set of experienced reward decrements *Δr* and the expected decrement 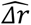. This series of calculations of differences in entropy before and after choice yields a smoothly declining function that represents the information participants gathered about the size of the change in reward from staying in a patch.

The model contained 10 parameters. As with the other models, this model contained a catch probability term for leaving patches on the first trial in patch as well as a side bias term to capture any choice biases due to the side of the screen that a target appeared on. In addition, the model contained a parameter for an intercept, the reward, the information, and the interaction of reward and information. After the first choice in a patch, leave probability was modeled as

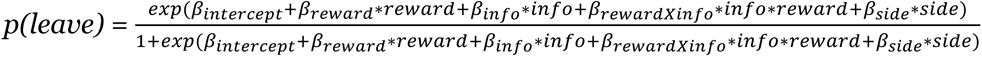

where *side* was coded as ‘1’ for left side and ‘0’ for right, *information* was the difference in entropy described above, and *reward* was one of the three currencies tested in all of the above models: reward for the current trial, time for the current trial, or reward rate for the current trial. This equation models decisions as a simple logistic transformation of those variables. Stay probabilities were computed as 1 - *p(leave)*. Because this model computes leave and stay probabilities using both *information* and *reward*, the model computes the probability of a decision using a mixture of currencies.

The mixture-of-currencies model, then, had 10 β parameters: side bias β_s_; one each of decision intercept β_0_, the reward parameter β_reward_, the information parameter β_info_, and the interaction parameter β_rewardXinfo_ for both patch types; and first trial leave catch probability β_1_.

#### Optimal Foraging Models

Optimal foraging theory (Stephens and Krebs 1986), a branch of formal ecology, explains foraging behavior in terms of mathematical models of behavior derived from biology, economics, and other disciplines. These models assess foraging decisions on the basis of reward rates. An exemplary model is the marginal value theorem (MVT) from Charnov (Charnov 1976), which dictates patch leaving decisions on the basis of a comparison of instantaneous to average reward rates. The MVT instructs foragers to depart depleting patches when the instantaneous reward rate falls below that average. The MVT model is a simple but powerful description of foraging behavior. Nonetheless, many species violate the predictions of the MVT, as discussed in the introduction. Inspired by the MVT, we fit several foraging models to participants’ choices.

##### 1. Learning MVT

The first model inspired by optimal foraging theory that we fit was adapted from Constantino and Daw (2015). Like the other models, choices were made by using a softmax equation

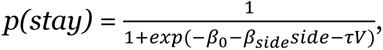

and probability of leaving *p(leave)* was

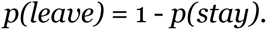

The variable for the side were defined as previously. The value *V* was computed as follows. A single patch value *V_patch_* was cached and on every choice, *V* was computed from the expected change in patch value, differently for each currency. For rewards,

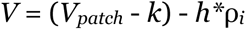

for the expected drop in reward *k*, handling time *h* equal to the median response time plus 1 for the time to initiate a trial, and *i*^th^ timeout type reward rate ρ*_i_*. The product of the handling time and reward rate amount to an expectation for the current trial’s reward. For reward rate,

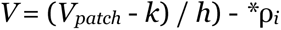

for the same definition of the variables as for reward. Subtracting the expected drop in reward from the patch value and dividing by the handling time amounts to an estimate of the current trial’s reward rate. Finally, for time,

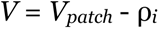

where ρ*_i_* is now the average time in patch of timeout type *i*. The value *V_patch_* was initialized at 7 for reward and reward rate and 0 for time, and then updated to the most recently received reward for reward and reward rate currencies and for the latest time in patch for time. These values were both set to 0 after leaving a patch. Each of these variables was updated over the course of the session separately for the 1 sec and 5 sec timeout blocks. For reward and reward rate, k was updated as

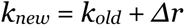

for single reward decrement *Δr*, and ρ*_i_* was updated as

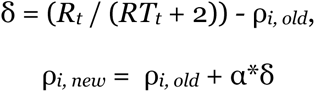

for learning rate α and trial *t*, recognizable as the basic reinforcement learning update rule. Time was updated similarly, with

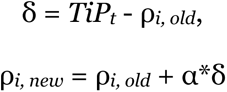

for time in patch *TiP_t_* on trial *t*.

The learning MVT model, then, had five β parameters: the decision noise τ; decision intercept β_0_; side bias β_s_; first trial leave catch probability β_1_; and the learning rate α.

##### 2. Learning Forager

We fit a second model inspired by optimal foraging theory, the learning forager model (McNamara and Houston 1985). Choices were made by using a softmax equation

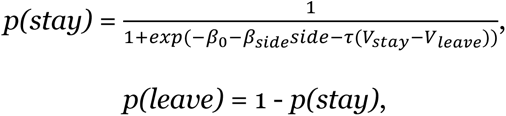

with the same variable and parameter definitions as the other models but for the value terms. The values were computed as follows. The initial rewards *R_0_* = 1 and the initial times *T_0_* = 1. For the reward rate currency, a running session reward rate

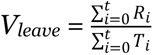

was compared to the current patch’s reward rate, the sum of the current patch’s rewards divided by the current patch’s total time in patch, for *V_stay_*. For reward, *V_leave_* was the sum of the session’s rewards and *V_stay_* the sum of the patch’s rewards, and for time, *V_leave_* was the sum of the session’s time in patch and *V_stay_* the patch’s time in patch; for both, the value of leaving was divided by the number of patches. Separate *V_leave_* values were computed for 1 sec and 5 sec timeout blocks.

The learning forager model had only four β parameters: the decision noise τ; decision intercept β_0_; side bias β_s_; and first trial leave catch probability β_1_.

##### 3. Recency Learning Forager

Finally, we fit a third optimal foraging model, a variant of the foraging learner model, the recency learning forager that weights the more recent outcomes in computing the value of staying or leaving (McNamara and Houston 1985).

Choices were again made by using a softmax equation

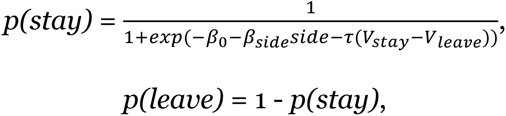

with the same variable and parameter definitions as the other models but for the value terms. For the reward rate currency, the value of leaving *V_leave_* on trial *t* was

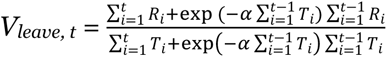

where α is a weight greater than 0. *R_0_* and *T_0_* were both set to 0. *V_stay_* equaled the sum of the current patch’s rewards divided by the total time in the current patch. For rewards, *V_leave_* was the sum of the session’s rewards divided by the number of patches, and for time, *V_leave_* was the total time in all patches divided by the number of patches. For the reward rate currency, *V_stay_* = the current patch’s reward rate, for rewards, *V_stay_* = the sum of the current patch’s rewards, and for time, *V_stay_* = the total time in the current patch. Separate *V_leave_* variables were computed for the 1 sec and 5 sec timeout blocks.

The recency learning forager model had five β parameters: the decision noise τ; decision intercept β_0_; side bias β_s_; first trial leave catch probability β_1_; and the recency weight α.

#### Monkey Model Fits

Monkeys were fit using the same set of 27 models as the humans. There were some minor adjustments to apply each model, which we briefly list here:

- Reinforcement learning models: the exact same models were used as the humans, except only a single set of Q-values was updated for patch states instead of a set for each timeout.
- Structure learning models: the exact same models were used as the humans.
- Mixture-of-currencies model: because the monkeys’ timeouts varied from 0.5 to 10.5 seconds, it was not possible to have separate parameters for each type of timeout. Instead, a single parameter was used for the intercept, reward, information and interaction of reward and information, yielding six parameters.
- Optimal foraging models: the exact same models were used as the humans, except a single leave value was used for all patches instead of one for each timeout.

## 6. Supplement

**Supplemental Table 1.** Model fit results for BIC. *: best-fit model overall. See methods for descriptions of each model.

| **Model Name:**<br>**currency** | **BIC Score: mean (s.e.m.)** |
| --- | --- |
| Patch State RL:<br>reward | 111.17 (1.30) |
| Patch State RL: time | 126.06 (1.50) |
| Patch State RL:<br>reward rate | 115.93 (1.33) |
| Patch State TD-RL:<br>reward | 113.53 (1.30) |
| Patch State TD-RL:<br>time | 119.95 (1.65) |
| Patch State TD-RL:<br>reward rate | 114.96 (1.29) |
| R Learning: reward | 117.68 (1.63) |
| R Learning: time | 126.55 (1.49) |
| R Learning: reward<br>rate | 119.12 (1.54) |
| Structure Learning:<br>reward | 103.60 (1.40) |
| Structure Learning:<br>time | 201.89 (9.82) |
| Structure Learning:<br>reward rate | 103.68 (1.39) |
| Structure Learning<br>with Discount: reward | 156.31 (2.14) |
| Structure Learning<br>with Discount: time | 421.88 (16.11) |
| Structure Learning<br>with Discount: reward<br>rate | 164.34 (2.04) |
| Learning MVT:<br>reward | 111.50 (1.48) |
| Learning MVT: time | 111.80 (2.76) |
| Learning MVT:<br>reward rate | 118.40 (1.58) |
| Learning Forager:<br>reward | 5689.60 (147.69) |
| Learning Forager:<br>time | 3662.24 (73.96) |
| Learning Forager:<br>reward rate | 129.19 (1.83) |
| Recency Learning<br>Forager: reward | 1139.92 (20.61) |
| Recency Learning<br>Forager: time | 816.78 (12.19) |
| Recency Learning<br>Forager: reward rate | 130.18 (1.85) |
| Mixture-of-currencies<br>Logistic: reward* | 98.48 (1.17) |
| Mixture-of-currencies<br>Logistic: time | 108.93 (1.35) |
| Mixture-of-currencies<br>Logistic: reward rate | 118.91 (1.34) |

**Supplemental Table 2.** Monkey model fit results. *: best-fit model overall. See methods for descriptions of each model. Each column reports the mean BIC score (s.e.m.) across sessions for that monkey.

| **Model Name: currency** | **BIC Score: M1** | **BIC Score: M2** |
| --- | --- | --- |
| Patch State RL: reward | 394.13 (30.88) | 596.20(45.69) |
| Patch State RL: time | 394.06 (31.07) | 595.74 (45.81) |
| Patch State RL: reward rate | 369.49 (32.73) | 594.74 (45.90) |
| Patch State TD-RL: reward | 347.97 (26.26) | 557.78 (45.28) |
| Patch State TD-RL: time | 364.41 (32.38) | 594.61 (42.72) |
| Patch State TD-RL: reward rate | 327.52 (19.12) | 566.64 (40.89) |
| R Learning: reward | 371.45 (29.16) | 559.99 (47.26) |
| R Learning: time | 371.06 (25.88) | 577.79 (43.02) |
| R Learning: reward rate | 352.47 (24.59) | 568.46 (46.58) |
| Structure Learning: reward | 358.72 (22.64) | 580.89 (43.74) |
| Structure Learning: time | 404.27 (30.36) | 596.14(45.31) |
| Structure Learning: reward rate | 366.61 (24.09) | 581.81 (43.85) |
| Structure Learning with Discount: reward | 682.66 (46.15) | 984.29 (80.64) |
| Structure Learning with Discount: time | 736.17 (56.36) | 1003.76 (96.93) |
| Structure Learning with<br>Discount: reward rate | 724.01 (65.97) | 1054.50 (77.18) |
| Mixture-of-currencies<br>Logistic: reward* | 87.53 (12.37) | 113.65 (38.93) |
| Mixture-of-currencies<br>Logistic: time | 302.05 (27.27) | 528.037 (39.65) |
| Mixture-of-currencies<br>Logistic: reward rate | 332.95 (26.23) | 575.00 (42.962) |
| Learning MVT: reward | 404.32 (30.35) | 597.38 (45.60) |
| Learning MVT: time | 402.37 (30.27) | 595.34 (45.48) |
| Learning MVT: reward rate | 303.59 (17.76) | 541.50 (43.44) |
| Learning Forager: reward | 427.64 (32.73) | 644.46 (48.97) |
| Learning Forager: time | 399.99 (30.41) | 591.82 (45.36) |
| Learning Forager: reward<br>rate | 1271.28 (195.60) | 2686.65 (233.19) |
| Recency Learning Forager:<br>reward | 433.52 (33.41) | 650.23 (49.23) |
| Recency Learning Forager:<br>time | 401.72 (30.57) | 593.72 (45.27) |
| Recency Learning Forager:<br>reward rate | 1264.67 (193.25) | 2677.43 (233.10) |

## Acknowledgements

The Wharton Behavioral Lab provided support in setting up and running the study. We are grateful to Emily M. Orengo and Richard Lee for their assistance during this project. We thank Nuwar Ahmed for help with data collection, Arjun Ramakrishnan for the original human task design, Benjamin Hayden for the original monkey task design, and Elizabeth Brannon for facilitation, oversight, and encouragement. The research was supported by R37-MH109728, R01-MH108627, R01-MH-118203, KA2019-105548, U01MH121260, UM1MH130981, R56MH122819, R56AG071023 (all to MLP), K99DA048748 to DLB, the Wharton Behavioral Lab and the Wharton Dean’s Research Fund (to VUL and MLP).

